# Comparative Genomics Reveals Genetic Factors Associated with Intraspecific Pigmentation Differences in Cassava

**DOI:** 10.64898/2026.09.18.752618

**Authors:** Moyosore Priscilla Shittu, Boas Pucker

## Abstract

Cassava produces starch-filled tubers which are a crucial food source for many people, especially in Africa. This plant species is famous for drought tolerance and resilience, but changing conditions due to climate change require genetic improvements to ensure sustainable cassava cultivation. In this context, a comprehensive understanding of the genetic basis responsible for a variety of traits is beneficial. Here, we present insights into the anthocyanin metabolism, responsible for coloration and protection of cassava structures during stress. Intraspecific variation in the transcriptional control of the anthocyanin biosynthesis was characterized through comparative genomics and phylogenetic analyses. A member of the R2R3-MYB transcription factor family, orthologous to crucial activators of anthocyanin biosynthesis in other plant species, was identified as a genetic hotspot for pigment accumulation capacity between cultivars. Orthologs of this gene appeared absent from three cassava cultivars without pronounced anthocyanin pigmentation, while being present in cultivars known for anthocyanin formation. This discovery corroborates previous reports about the importance of transcriptional regulators in the intraspecific diversity of anthocyanin pigmentation.

## Introduction

*Manihot esculenta* (Cassava) is a vital staple crop that feeds over 500 million people (Otekunrin, 2024), particularly in sub-Saharan Africa. Cassava is processed into different food products and every part of the plant is either eaten or used to regrow a plant. Cassava serves as an essential source of carbohydrates for animal and human consumption. The ability to survive in arid regions is especially important, making cassava useful for commercial agriculture (Immanuel *et al*., 2024). With a 61% production rate, Africa has the highest cassava production percentage, which is the largest in the world and exceeds the continent’s maize production by a factor of two (Wosula *et al*., 2024; Otekunrin, 2024). This reflects the importance of cassava in everyday activities. To sustain cassava cultivation in a climate-changing environment and enhance nutritional value, breeding and genome-editing efforts are needed. Both approaches benefit from a comprehensive understanding of the genetics underlying important traits. Modern crop improvements rely on the availability of a genome sequence to inform molecular biology studies. The genome of several cassava cultivars has been sequenced (Qi *et al*., 2022; Landi *et al*., 2023; Thoben *et al*., 2025; Beier *et al*., 2025), thus providing a comprehensive genomic basis for comparative investigations.

Anthocyanin accumulation is a visual trait linked to a range of physiological functions and stress-response processes (Winkel-Shirley, 2001; Grünig *et al*., 2025; Choudhary *et al*., 2026b). Anthocyanins are plant specialized metabolites, produced through a branch of flavonoid biosynthesis and well-known for their colors (Winkel-Shirley, 2001; Grotewold, 2006; Grünig *et al*., 2025). Many of the blue, pink, red, and purple colors in vegetables and fruits, which play a vital role in plant propagation, ecophysiology, and plant defense mechanisms, are conferred by anthocyanins (Alappat & Alappat, 2020; Grünig *et al*., 2025; Muralidhar *et al*., 2026).

Anthocyanins are derived from the aromatic amino acid phenylalanine, which is processed through the general phenylpropanoid pathway into flavonoid biosynthesis (**Figure 1**). Structural genes of the flavonoid biosynthesis encode the enzymes chalcone synthase (CHS), chalcone isomerase (CHI), and flavanone 3-hydroxylase (F3H) that channel the substrate towards anthocyanin biosynthesis (Winkel-Shirley, 2001). A pragmatic definition of anthocyanin biosynthesis typically encompasses the enzyme-catalyzed reactions mediated by dihydroflavonol 4-reductase (DFR), anthocyanidin synthase/leucoanthocyanidin dioxygenase (ANS/LDOX), anthocyanin-related glutathione S-transferase (arGST), and UDP-dependent anthocyanidin 3-O-glycosyltransferase (UGT) (Choudhary *et al*., 2026b). DFR competes with the flavonol synthase (FLS) for substrate (Choudhary & Pucker, 2024). Leucoanthocyanidin reductase (LAR) and anthocyanidin reductase (ANR) catalyze the formation of proanthocyanidins and represent additional reactions competing with the formation of anthocyanins (Choudhary *et al*., 2026b). Synthesis of anthocyanins at the endoplasmic reticulum is followed by intracellular transport into the central vacuole, where the pigments accumulate (Grünig *et al*., 2025).

**Figure 1:**
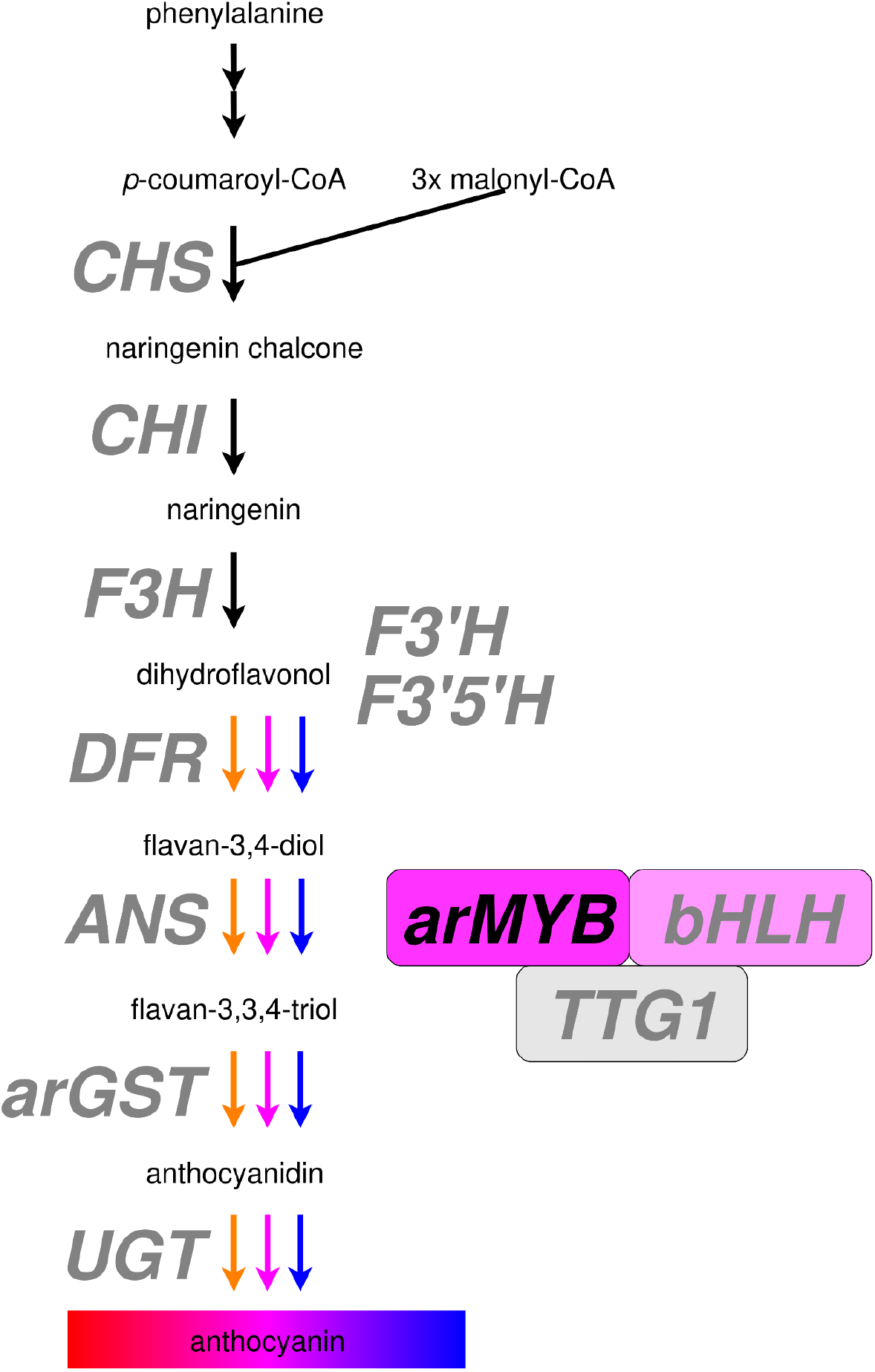
Simplified illustration of anthocyanin biosynthesis. CHS, chalcone synthase; CHI, chalcone isomerase; F3H, flavanone 3-hydroxylase; F3’H, flavonoid 3’-hydroxylase; F3’5’H, flavonoid 3’,5’-hydroxylase; DFR, dihydroflavonol 4-reductase; ANS, anthocyanidin synthase; arGST, anthocyanin-related glutathione S-transferase; UGT, UDP-dependent anthocyanidin 3-O-glycosyltransferase; arMYB, anthocyanin-related MYB transcription factor; bHLH, basic helix-loop-helix transcription factor; TTG1, TRANSPARENT TESTA GLABRA 1. This figure layout has been derived from (Horz *et al*., 2026).

Anthocyanin biosynthesis is controlled at the transcriptional level through the interaction of positive and negative regulators (Grünig *et al*., 2025). The most important activator of anthocyanin biosynthesis is the MBW complex (Gonzalez *et al*., 2008) formed by an R2R3-MYB transcription factor protein, a bHLH protein, and the WD40 protein TRANSPARENT TESTA GLABRA 1 (TTG1). This complex is involved in a range of plant functions, with specificity resulting from the integration of different MYB (myeloblastosis) and bHLH (basic helix-loop-helix) proteins (Ramsay & Glover, 2005) (Li, 2014; Xu *et al*., 2015). Members of the MYB subgroup 6 and subgroup 5 have been reported as anthocyanin-specific activators (Gonzalez *et al*., 2008; Marin-Recinos & Pucker, 2024; Khatun *et al*., 2025). Negative regulators counter the MBW complex’s activation. According to (LaFountain & Yuan, 2021), these repressors fall into distinct functional groups, with the MYB4 lineage representing one of the most important negative regulators of phenylpropanoid and flavonoid biosynthesis (Jin *et al*., 2000; Wang *et al*., 2020; Deng *et al*., 2021). Other MYB repressors such as FaMYB1 (Aharoni *et al*., 2001), MYBC2 (Cavallini *et al*., 2015), and MYBL2 (Dubos *et al*., 2008) are also negative regulators of the anthocyanin pathway. Anthocyanin biosynthesis has also been well studied in model species such as *Petunia hybrida, Antirrhinum majus*, and *Arabidopsis thaliana*, which were paramount in establishing the regulatory framework of the anthocyanin pathway (Goff *et al*., 1992; Quattrocchio *et al*., 1999; Gonzalez *et al*., 2008). Recent analyses have expanded this classical framework and revealed substantial evolutionary flexibility in anthocyanin biosynthesis (Khatun *et al*., 2025; Choudhary *et al*., 2026b,a). In cassava, the presence of these transcription factors has been documented (An *et al*., 2022), but these factors have not yet been investigated in depth with respect to anthocyanin regulation. Identifying and characterizing the transcriptional regulators of anthocyanin biosynthesis will provide a framework for crop improvement endeavours.

The aim of this study is to identify and characterize genes encoding anthocyanin biosynthesis-related proteins, including positive and negative transcriptional regulators in cassava to explain phenotypic variation in the species. The presence/absence of a single R2R3-MYB transcription factor was tightly coupled to the anthocyanin biosynthesis capacity of cassava accessions.

## Material and Methods

### Data collection

Cassava genomic and transcriptomic datasets were retrieved from the National Center for Biotechnology Information (NCBI; https://www.ncbi.nlm.nih.gov/) (Goldfarb *et al*., 2025), Phytozome (Goodstein *et al*., 2012), and CASSAVASTORE hosted on the PlaBiPD platform (Beier *et al*., 2025). A total of 23 *Manihot* genome assemblies representing diverse cultivars and breeding lines were collected alongside closely related outgroups. GCA_000737105.1 (W14) was previously classified as a wild cassava ancestor (*M. esculenta ssp. flabellifolia*) and was treated as an unresolved *Manihot* outgroup closely related to *M. glaziovii*, in line with corrected genomic classifications (Lyons *et al*., 2022) and GCA_965364665.1 (Cassava wild type). An analysis with BUSCO v6.0.0 (Tegenfeldt *et al*., 2025) was conducted in protein mode using the embryophyta_odb12 lineage dataset to assess the completeness of the annotated polypeptide sequences for each cultivar.

### Annotation of unannotated genome sequences

Out of the 23 retrieved cassava genome assemblies, 11 assemblies lacked a structural annotation and were therefore annotated using GeMoMa v1.9 (Keilwagen *et al*., 2019). The two closely related species *Jatropha curcas* (GCA_000696525.1) (Zhang *et al*., 2014) and *Hevea brasiliensis* (GCA_000696525.1) (Cheng *et al*., 2023), served as hints for gene prediction (Additional File A). AGAT (Dainat *et al*., 2024) was applied for cleaning and generating coding sequence (CDS) and polypeptide sequence (PEP) files for the genome sequences without publicly available annotations.

### Identification of potential repressors

A literature search was carried out to identify characterized anthocyanin repressors from model and non-model plants (Additional File B). The classification framework described by LaFountain & Yuan (2021), was used to guide assignment of sequences to known repressor types. Transcriptional repressors associated with the anthocyanin biosynthesis include MYB4, FaMYB1, MYBC2, and MYBL2. Orthologous proteins were identified based on their position in a gene tree following the collection of the sequence of the best BLAST hit per bait based on collect_best_BLAST_hits.py (Pucker & Iorizzo, 2023), executed with the manually compiled bait collection (Additional File B), to identify repressors in cassava and a set of 260 plant species (Additional File C). The resulting collection of putative repressor sequences was filtered through the construction of gene family trees for each putative repressor. Reference genome assembly (cultivar AM560-2, v8) was used for gene identification, and it follows the naming convention Manes.XXGXXXXXX to identify the putative cassava repressors, where “Manes” denotes a *Manihot esculenta locus, the leading* digits indicate the chromosome, and the trailing digits indicate the gene identifier on that chromosome. In brief, for each putative repressor family homologous sequences in a set were aligned with MAFFT v7.525 (L-INS-i algorithm) (Katoh & Standley, 2013), alignments were trimmed pxclsq from the PHYX toolkit (-p 0.1) (Brown *et al*., 2017), phylogenies were inferred by IQ-TREE3 v3.1.1 (Wong *et al*., 2026), and the visualization was achieved in iTOL (Letunic & Bork, 2024).

### Structural gene identification in the anthocyanin pathway

Structural genes in the phenylpropanoid and flavonoid pathways (*CHS, CHI, F3H, F3’H, DFR, ANS, arGST, UGT*) were identified using KIPEs v3.2.6 (Rempel *et al*., 2023), which allows for the identification of candidate sequences based on orthology to characterized baits and inspection of functionally important amino acid residues in all candidates. KIPEs was run across the polypeptide sequence sets of 260 plant species (Additional File C), including angiosperm and gymnosperm species, to enable an assessment of conservation patterns between the cassava cultivars and other plant species.

### Identification of key transcription factors

To identify transcription factors associated with anthocyanin regulation, two specialized annotation tools (MYB_annotator and bHLH_annotator) were applied. MYB_annotator v1.0.3 (Pucker, 2022) was used to characterize all R2R3-MYB genes, while the bHLH_annotator v0.2 (Thoben & Pucker, 2023) was used to annotate basic helix-loop-helix transcription factors. Important anthocyanin biosynthesis regulators like anthocyanin-related MYB (arMYB) and the bHLH protein TRANSPARENT TESTA 8 (TT8) were extracted from the MYB_annotator and bHLH_annotator results, respectively. The identification of TTG1 was based on an orthology assessment based on a previously reported TTG1 protein sequence collection (Choudhary *et al*., 2026a).

### Phylogenetic analysis and microsynteny analysis

To infer evolutionary relationships among repressing and activating transcription factors, polypeptide sequences were aligned using MAFFT v7.505 (Katoh & Standley, 2013). Alignments were cleaned with pxclsq from the PHYX v1.3.2 toolkit (Brown *et al*., 2017) with a column-filtering threshold of -p 0.1 to remove column with more than 90% gaps, and maximum likelihood phylogenies were inferred by IQ-TREE v3.1.0 with 1000 bootstrap replicates (Wong *et al*., 2026). For the arMYB tree **(Figure 2)**, we selected the JTTDCMUT+R5 model based on the Bayesian Information Criterion (BIC). For the repressor MYB tree (Additional File H), we selected JTT+R10 based on the Bayesian Information Criterion (BIC). Gene trees were inspected using iTOL v6 (Letunic & Bork, 2024). We used these trees to classify cassava repressors into established groups such as MYB4, FaMYB1, MYBC2, and MYBL2. Microsynteny analysis was conducted using JCVI/MCscan v1.6.7 (Tang *et al*., 2024) to compare genomic loci of interest between the 23 cassava cultivars.

**Figure 2:**
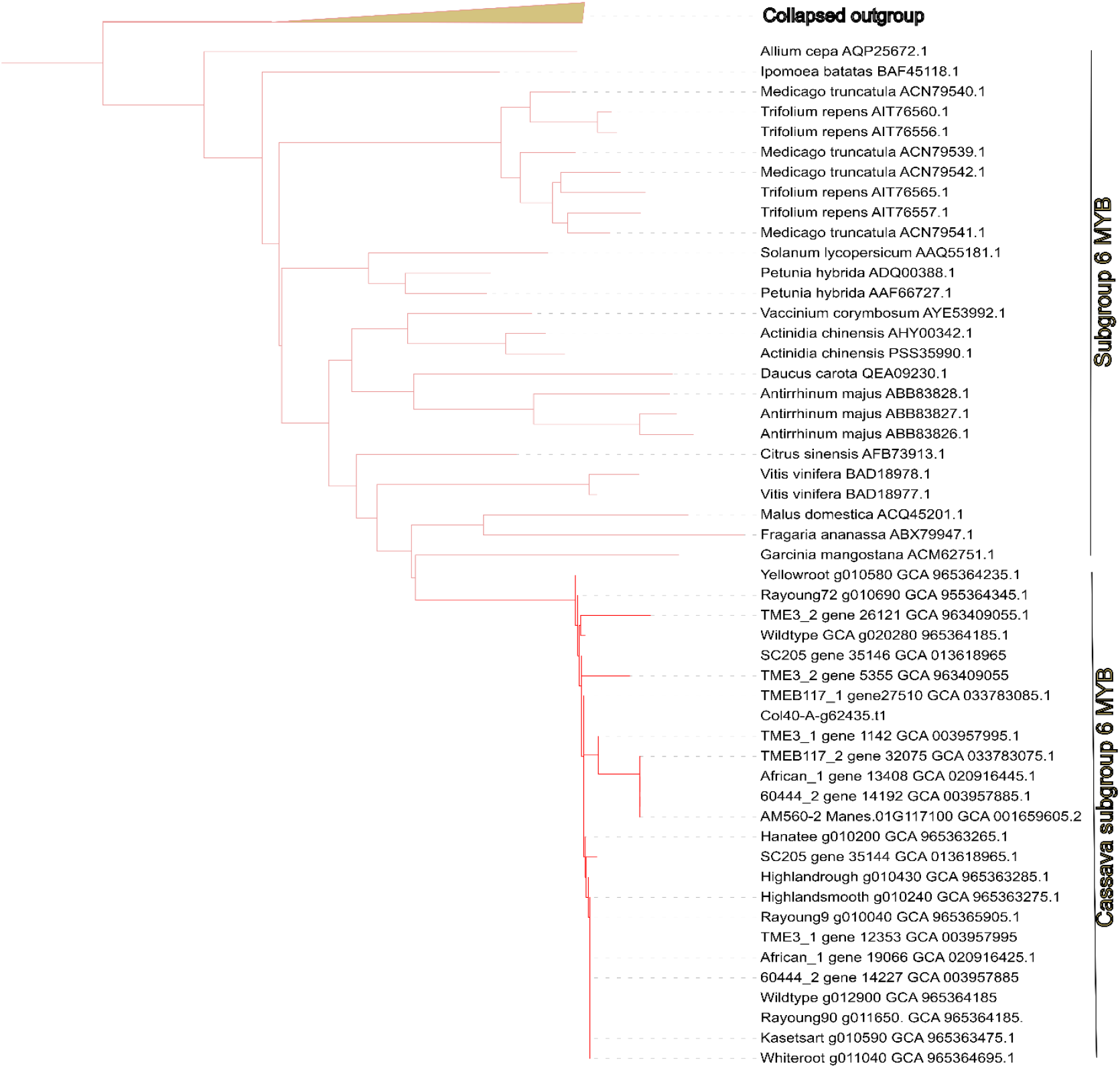
Subgroup 6 (SG6) R2R3-MYB transcription factors tree associated with anthocyanin biosynthesis activation (arMYB) in a lighter shade of red. The presence of numerous cassava cultivar sequences in a darker shade of red shows that orthologous MYBs are generally detectable. No sequences from the three analyzed cultivars W14, KU50, and 60444 were detected in this arMYB clade, suggesting absence.

**Figure 3:**
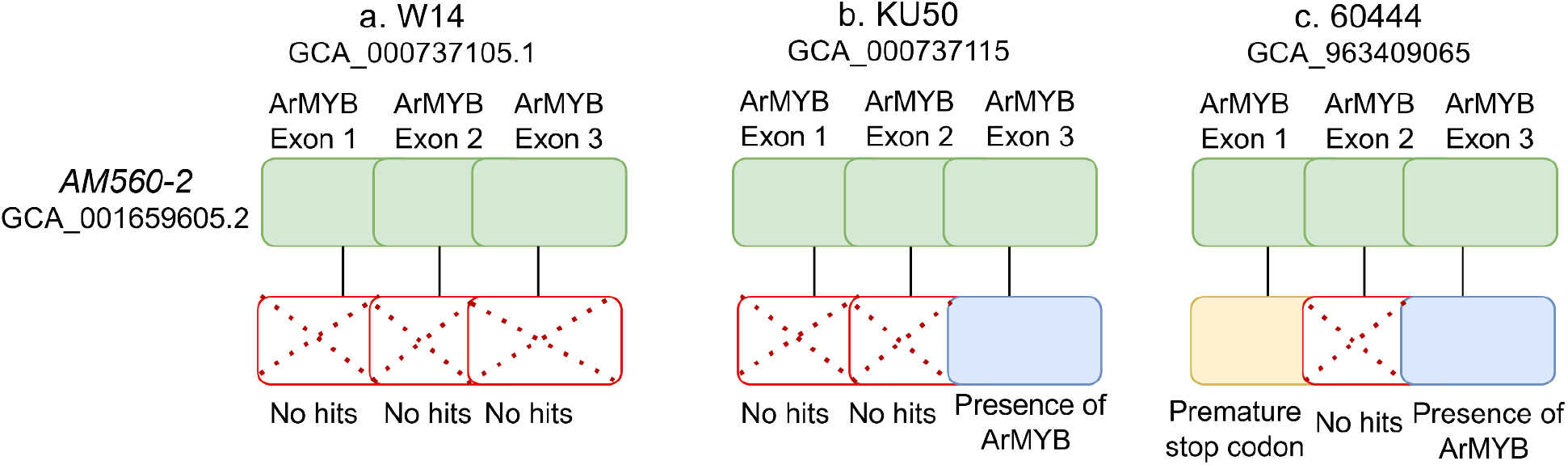
Schematic representation of the three arMYB exons in the AM560-2 reference genome (green, top) and corresponding tBLASTn evidence recovered from the genome assemblies of W14 (GCA_000737105.1), KU50 (GCA_000737115.1), and 60444 (GCA_963409065.1) (bottom). Blue boxes indicate exons recovered as high-confidence matches to the reference; red hatched boxes indicate exons for which no true ortholog was detected; the gold coloured box in 60444 indicates an exon identified up to a premature stop codon. No exon returned a confident match in W14 (a). In KU50, only exon 3 was recovered (b).

### Direct sequence comparison and structural integrity assessment of arMYB in cassava cultivars

To assess the presence and structural integrity of arMYB (MYB113; Manes.01G117100 in the *Manihot esculenta* v8 reference) in cultivars W14 (GCA_000737105.1), KU50 (GCA_000737115.1), and 60444 (GCA_963409065.1), direct sequence comparisons was carried out using tblastn (BLAST v2.17.0) (Altschul *et al*., 1990) against raw genome assemblies, independent of gene annotation. Whole-protein searches used an E-value threshold of 1 × 10^−5^; to resolve ambiguous hits, the arMYB protein was divided into its three exons (residues 1-39, 40-82, and 83-235) and searched individually (E-value ≤ 1). A hit was considered a true ortholog only if it exceeded 85% identity over ≥25 residues. When BLAST hits for arMYB and its flanking genes mapped to the same scaffold, the region was extracted using samtools faidx (v1.19.2) (Danecek *et al*., 2021) and translated in the relevant reading frame using seqkit (v2.3.0) (Shen *et al*., 2024) to verify reading-frame continuity and identify premature stop codons. Method reliability was validated using two chromosome-level assemblies (Kasetsart50 and Rayong90), which correctly recovered all three arMYB exons at high identity (91.5-100%) in the expected order.

## Results and Discussion

### Structural genes of the anthocyanin biosynthesis are conserved across cassava

Structural genes involved in the anthocyanin biosynthetic pathway were highly conserved across all 23 cassava cultivars (Additional File D, Additional File E). The core anthocyanin pathway enzymes for the production of anthocyanin were identified, including CHS, CHI, F3H, DFR, ANS, arGST, and UGT. The presence of these genes across assemblies of the different cassava cultivars indicates that all possess the full enzymatic capacity to synthesize anthocyanins. This is consistent with its ability to produce pigmentation in leaves, stems, flowers, and storage roots as previously reported (Luo *et al*., 2023).

Among the anthocyanin biosynthesis enzymes, DFR was the only one that showed a notable amino acid substitution from the expected residues. All cassava DFR sequences showed a single residue variation at position 227, where histidine (H) replaced the conserved asparagine (N) or glutamine (Q) typically found in other angiosperms (Additional File F). All cassava cultivars studied here showed only a single DFR copy. This substitution is not specific to cassava, but was also consistently observed across representative Euphorbiaceae species, namely *Jatropha curcas, Euphorbia lathyris, Mercurialis annua, Ricinus communis, Hevea brasiliensis, and Linum usitatissimum* (Linaceae). This suggests that the amino acid substitution in DFR might be order-specific for the Malpighiales. Numerous reports about the presence of anthocyanins in Euphorbiaceae (Fu *et al*., 2022; Suresh *et al*., 2026) strongly suggest that this DFR variant must be functional. The generally high level of conservation across the anthocyanin biosynthesis enzymes underscores the evolutionary stability of the pathway in cassava and suggests that variation in pigmentation among cultivars is unlikely to arise from loss or disruption of structural gene function. It appears more likely that regulatory differences such as transcription factor activity or repressor diversification are responsible.

### MBW activators show regulatory variation

As the central regulator of anthocyanin biosynthesis, the MBW complex components were explored across the cassava cultivars. Members of subgroup 6 of the R2R3-MYB transcription factor family confer anthocyanin specificity to the MBW complex (Gonzalez *et al*., 2008; Marin-Recinos & Pucker, 2024). Numerous cases of intraspecific anthocyanin variation suggest that this MYB is the decisive factor in controlling anthocyanin production (Marin-Recinos & Pucker, 2024). *A. thaliana* harbours four close paralogs of this anthocyanin-related gene, namely MYB75/PAP1, MYB90/PAP2, MYB113, and MYB114 (Stracke *et al*., 2001). Only a single orthologous gene, highly similar to MYB113, was identified in cassava. The absence of the other three paralogs suggests that the gene duplication event giving rise to these four genes occurred after the divergence of *Arabidopsis* and cassava (Lynch & Conery, 2000). Therefore, the cassava MYB113-like gene is the most likely ortholog of the ancestral subgroup 6 MYB gene and was used as the representative member of subgroup 6 for comparative analysis in this study. This supports a clear 1:1 orthologous relationship between cassava and the ancestral lineage of Arabidopsis subgroup 6 MYBs, with MYB113 serving as the best functional and evolutionary proxy. A comprehensive gene tree of this anthocyanin MYB and orthologs in numerous plant species suggested the absence of the anthocyanin biosynthesis-activating subgroup 6 MYBs in the cassava cultivars W14, KU50, and 60444 (**Figure 2**).

The absence of the other three paralogs suggests that the gene duplication event giving rise to these four genes occurred after the divergence of Arabidopsis and cassava. Therefore, the cassava MYB113-like gene is the most likely ortholog of the ancestral subgroup 6 MYB gene and was used as the reference for comparative analysis. This supports a clear 1:1 orthologous relationship between cassava and the ancestral lineage of Arabidopsis subgroup 6 MYBs, with MYB113 serving as the best functional and evolutionary proxy.

TT8, the principal bHLH partner of MYB activators in the MBW complex, showed copy number variations across the 23 cassava cultivars, with numbers ranging from 1 to 8. These duplications may drive differences in regulatory strength or tissue-specific activation of anthocyanin biosynthesis. In contrast, GL3, another bHLH capable of participating in MBW formation, has not been detected in four cultivars. Because TT8 is present in all cultivars, the absence of another bHLH protein is unlikely to compromise MBW functionality.

### Genomic evidence for *arMYB* loss and pseudogenization

To explore whether the absence of arMYB in W14, KU50, and 60444 is due to a lack of detection or gene loss events, direct sequence-level evidence in each cultivar’s genome assembly was examined. Microsynteny analysis was also attempted around this locus but proved inconclusive, as annotation gene identifiers do not reflect true positional order and several assemblies are too fragmented to reliably reconstruct local gene order; we therefore relied on direct sequence comparison instead. We validated this approach using two chromosome-level reference assemblies (Kasetsart50 and Rayong90), in which all arMYB exons were recovered at high identity (91.5-100%) and in the expected order, confirming that the method reliably detects an intact ortholog when present. Although ultimate proof of gene loss is theoretically impossible, this method provides the strongest possible support for such a scenario; convergent evidence from whole-gene, exon-level, and flanking-region sequence searches strongly supports genuine structural disruption of arMYB in the three cultivars examined below.

In W14, the flanking genes of *arMYB* were detectable but located on different scaffolds, preventing identification of a clear continuous syntenic region surrounding the expected *arMYB* locus. No exon of arMYB itself showed a true match anywhere in the genome sequence, with the best BLAST hits reaching below 74% identity. We speculate that a structural variation could have resulted in the loss of this MYB gene.

In KU50, only exon 3 of arMYB is detectable with 91.5% identity, while exons 1 and 2, which together encode the R2R3 DNA-binding domain, were undetectable, indicating a partial deletion that removes the functionally essential region of the protein, thus confirming the absence of a functional arMYB gene.

In 60444, only relics of arMYB involving the start of exon 1 were recovered for its first 28 of 39 residues before a premature stop codon; exon 2 was undetectable, and exon 3 was intact (90.2% identity). If expressed, the resulting gene product from this locus would therefore have only approximately 28 amino acids due to a premature stop codon in exon 1, consistent with pseudogenization rather than complete deletion.

Given the established role of arMYB as a central activator of anthocyanin biosynthesis, its loss or disruption in these cultivars would explain the variation in pigmentation-related traits. This observation aligns with previous studies that have identified orthologs of this R2R3-MYB as the causal factor for anthocyanin loss at the species-level (Marin-Recinos & Pucker, 2024).

In 60444, exon 1 was recovered up to a stop codon, exon 2 was undetected, and exon 3 was intact (c). Diagrams are schematic and not drawn to scale.

### Investigation of four distinct anthocyanin repressor clades

Four anthocyanin associated repressors (MYB4, FaMYB1, MYBC2, and MYBL2) have been described previously (LaFountain & Yuan, 2021). To identify putative anthocyanin repressors in cassava, we performed phylogenetic clustering of cassava polypeptide sequences with experimentally validated repressors from diverse plant species. Only cassava sequences that clustered within well-supported clades containing characterized repressor sequences and *Amborella trichopoda* sequences at the base, indicating ancestral position, were considered orthologous. This phylogenetic analysis revealed that cassava repressors clustered into several well-supported clades corresponding to the major anthocyanin-associated MYB repressor families (MYB4, FaMYB1, MYBC2, MYBL2). FaMYB1 and MYBC2 clustered together, firmly indicating they are close relatives to each other and not distant, with no evidence for a distinct MYBL2 clade in cassava.

#### MYB4 repressors

One cassava MYB4 candidate clustered with a characterized MYB4 repressor (CAD98762.1, *Populus tremula* x *Populus tremuloides*, (Dubos *et al*., 2008)), including the previously characterized *Arabidopsis thaliana* sequence. The putative MYB4 repressor in cassava is *Manes*.*04G074900* (**Figure 4**). Functionally, MYB4-type transcription factors operate as key negative regulators across general phenylpropanoid and flavonoid metabolism (LaFountain & Yuan, 2021). They modulate metabolic flux either through direct transcriptional repression by binding to the promoters of downstream structural genes (such as *CHS, DFR, ANS*, and *UFGT*) or by establishing feedback regulatory loops with activator MYBs to prevent excessive pigment deposition and maintain cellular homeostasis (Cavallini *et al*., 2015; LaFountain & Yuan, 2021; Chen *et al*., 2026).

**Figure 4:**
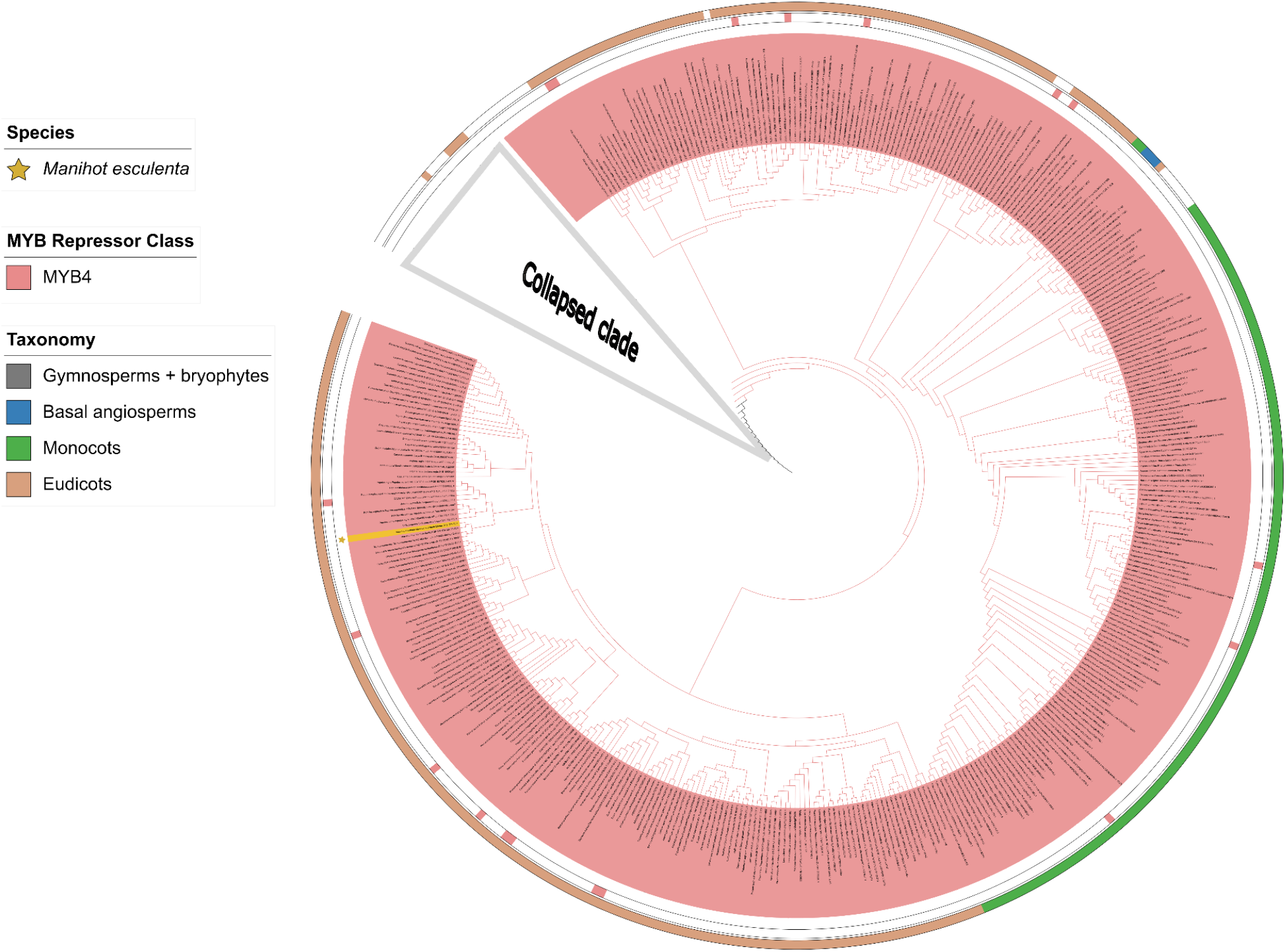
MYB4 clade phylogenetic tree. Sequences assigned to the MYB4 clade are highlighted with a red background. Color markings around the tree indicate the taxonomic lineages to which the neighbouring sequences belong. Markings between the outer circles and the sequence names indicate the positions of previously characterized MYB sequences. The putative cassava repressor is highlighted in gold. A tree figure with high resolution is available as Additional file I.

#### FaMYB1, MYBC2, and MYBL2 repressors

Phylogenetic analysis identified three candidate repressor genes in cassava within the FaMYB1/MYBC2/MYBL2 clade. Among these, *Manes*.*17G075600* is a cassava candidate that clustered directly alongside characterized *Prunus persica* FaMYB1 orthologs (ALO81021, ALO81022, KT159234) representing the primary conserved *FaMYB1* lineage in cassava. In contrast, two cassava genes, *Manes*.*02G074300* and *Manes*.*01G115400*, formed a distinct and cassava-specific clade. Their closest relatives in the tree were uncharacterized MYB sequences from *Ricinus communis* and *Vernicia fordii*, both Euphorbiaceae, supporting their placement in the tree. The characterized FaMYB1 ortholog AJI76863.1 (*Populus tremula* × *P. tremuloides*; (Yoshida *et al*., 2015)) was the nearest characterized reference to this clade for *Manes*.*02G074300* and *Manes*.*01G115400* in the clade (**Figure 5)**. Notably, there are some inconsistencies among the reference sequences. For example, closely related cotton (*Gossypium*) paralogs have independently been annotated in the literature as FaMYB1 (AAN28286.1, *Gossypium hirsutum* (Cao *et al*., 2017)) and MYBL2 (ABX26107.1, *Gossypium hirsutum* (Dubos *et al*., 2008)) despite their small phylogenetic distance. This suggests the boundaries between the FaMYB1, MYBC2, and MYBL2 designations are not phylogenetically discrete even among well-studied reference sequences, which weakens the case for assigning cassava’s candidates definitively to any one of these three named classes.

**Figure 5:**
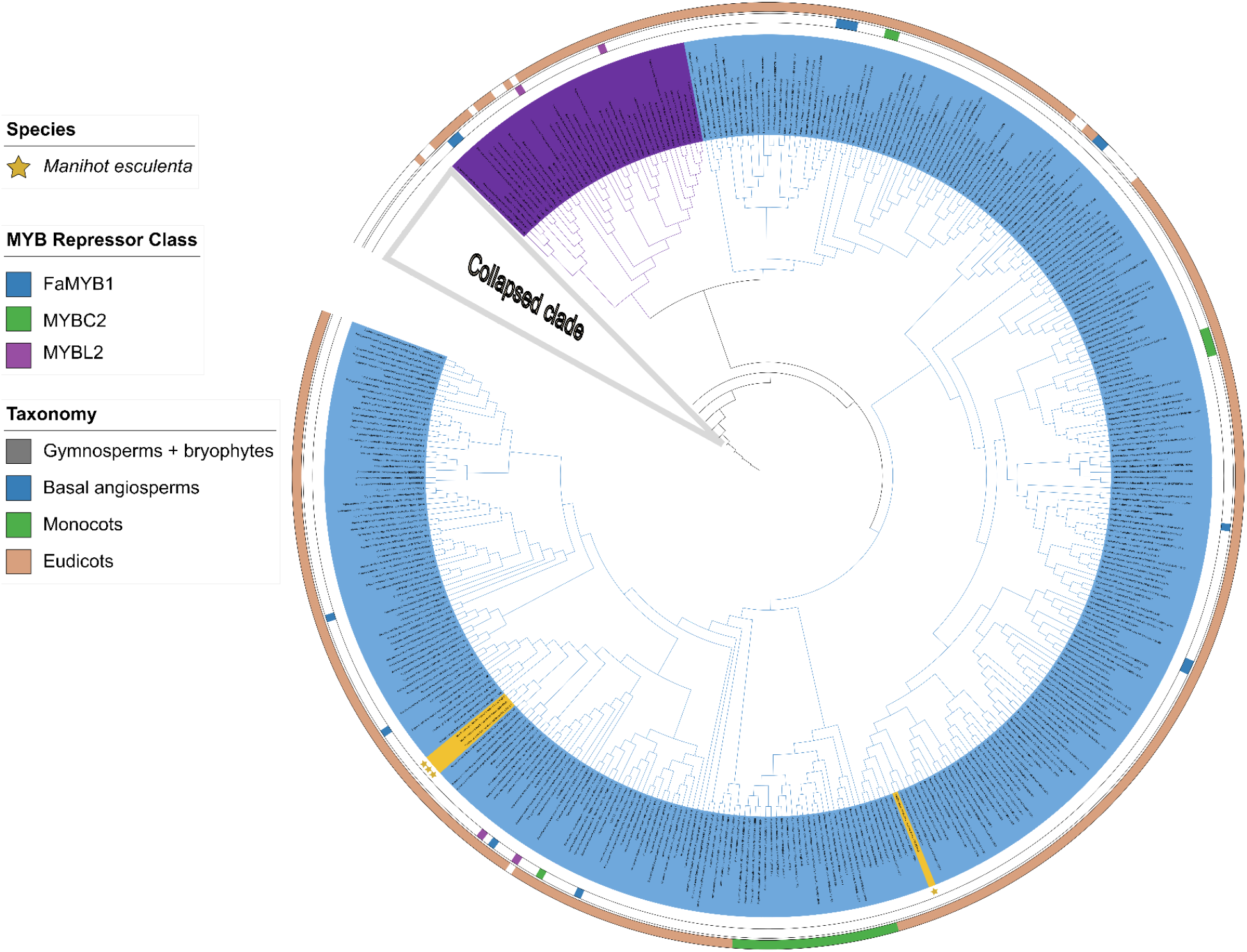
Phylogenetic tree of FaMYB1, MYBC2, and MYBL2. Sequences assigned to the FaMYB1 clade are highlighted with a blue background; sequences assigned to the MYBL2 clade are highlighted with a purple background. Color markings around the tree indicate the taxonomic lineages of the neighbouring sequences. Markings between the outer circles and the sequence names indicate the positions of previously characterized MYB sequences. The classification of MYB genes into the FaMYB1, MYBC2, and MYBL2 lineages remains inconsistent across the literature, with varying criteria and nomenclature complicating comparative assignments. This lack of consensus, rather than a fundamental ambiguity in phylogenetic resolution, suggests that assigning cassava MYB candidates to these three classes based solely on tree topology may be premature. A more cautious, integrative approach that considers sequence context, synteny, and functional evidence is therefore advisable to ensure accurate and reproducible classification. A high-resolution tree figure is available as Additional file J.

Nevertheless, members across these repressor classes share conserved negative-regulatory roles either directly by repressing late biosynthetic steps, as established for grapevine MYBC2 (Xie et al., 2020), or by disrupting MBW complex assembly, as observed for MYBL2 and FaMYB1 orthologs in other eudicots (Paolocci *et al*., 2011; Zhang *et al*., 2026). While the presence of this clade reflects broad evolutionary conservation across plants, the specific duplication of *Manes*.*02G074300* and *Manes*.*01G115400* in cassava suggests these two genes may have evolved specialized or distinct regulatory roles. This gene duplication could allow them to separately control storage root pigmentation, stress responses, or cultivar-specific color variation, though functional validation will be required to confirm their precise activities. This duplication pattern has been observed in other plant MYB families, such as the subgroup 7 R2R3-MYBs in *Arabidopsis thaliana*, where gene duplication led to subfunctionalization in flavonol regulation (Stracke *et al*., 2001, 2007).

The MBW activators, structural genes, and repressor families can be integrated into a unified regulatory model of cassava anthocyanin biosynthesis (**Figure 6**).

**Figure 6:**
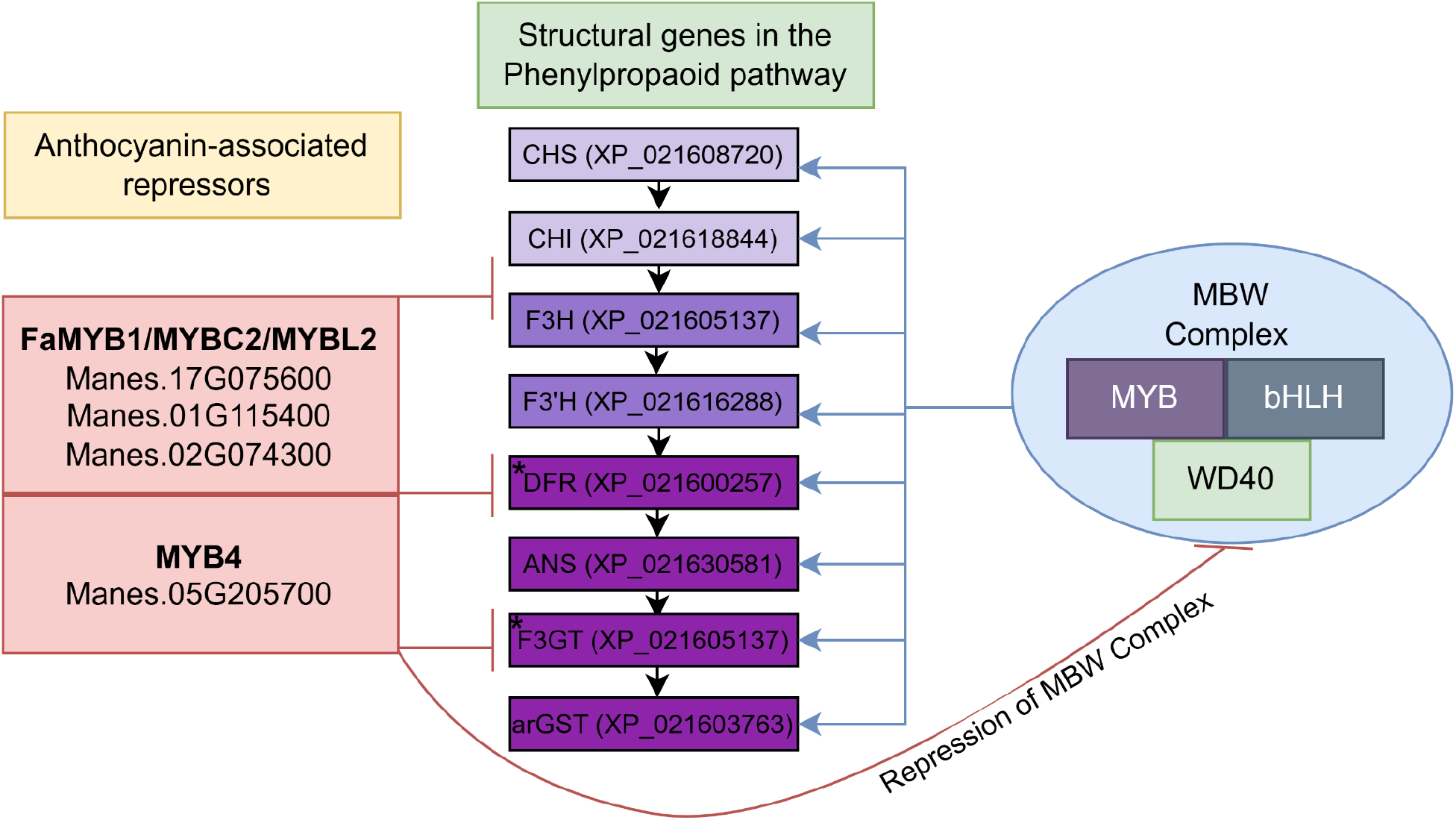
Proposed model of the Phenylpropanoid pathway and anthocyanin biosynthesis regulation in cassava. The MBW Complex (arMYB, TT8, TTG1) activates the late biosynthesis genes (DFR, ANS, arGST, UGT) responsible for anthocyanin biosynthesis. MYB repressors (MYB4, FaMYB1, MYBC2) are transcriptionally induced and provide feedback inhibition of the MBW complex, forming a multilayered regulatory loop. DFR and F3GT have asterisks because they are missing an amino acid residue in all *Manihot* cultivars analyzed.

## Data availability

All data sets underlying this study are publicly available. Specific locations of datasets are listed in the supplementary files. Novel structural annotations of cassava genome sequences are available via bonndata: https://doi.org/10.60507/FK2/IHEIMY.

## Acknowledgements

This work was supported by the de.NBI Cloud within the German Network for Bioinformatics Infrastructure (de.NBI) and ELIXIR-DE (Forschungszentrum Jülich and W-de.NBI-001, W-de.NBI-004, W-de.NBI-008, W-de.NBI-010, W-de.NBI-013, W-de.NBI-014, W-de.NBI-016, W-de.NBI-022). We thank all members of the Plant Biotechnology and Bioinformatics research group for their discussion and support. We acknowledge support from Project DEAL and the University of Bonn for open access publication. MPS is grateful for a Jeff Schell Fellowship for Agricultural Science provided by the Bayer Foundation.

## Author contributions

MPS and BP designed the study. MPS conducted the bioinformatic analyses. MPS and BP wrote the manuscript. All authors reviewed the final version of the manuscript and consented to its submission.

## Additional Files

Additional File A: Datasets used as hints for structural annotation.

Additional File B: Identified anthocyanin-associated repressor bait sequences

Additional File C: Names of all 260 species represented in the sequence collection for repressor identification

Additional File D: All anthocyanin biosynthesis gene IDs from the KIPEs result

Additional File E: Cassava cultivar IDs used for the KIPEs analysis

Additional File F: Alignment of DFR sequences of different cassava cultivars

Additional File G: DFR alignment showing the DFR substitution amongst the Euphorbiaceae

Additional File H: Full repressor tree

Additional File I: MYB4 repressor tree

Additional File J: FaMYB1, MYBC2, and MYBL2 repressor tree

