## Supplementary figures and images for "Comparative Genomics Reveals Genetic Factors Associated with Intraspecific Pigmentation Differences in Cassava"

### Additional File H

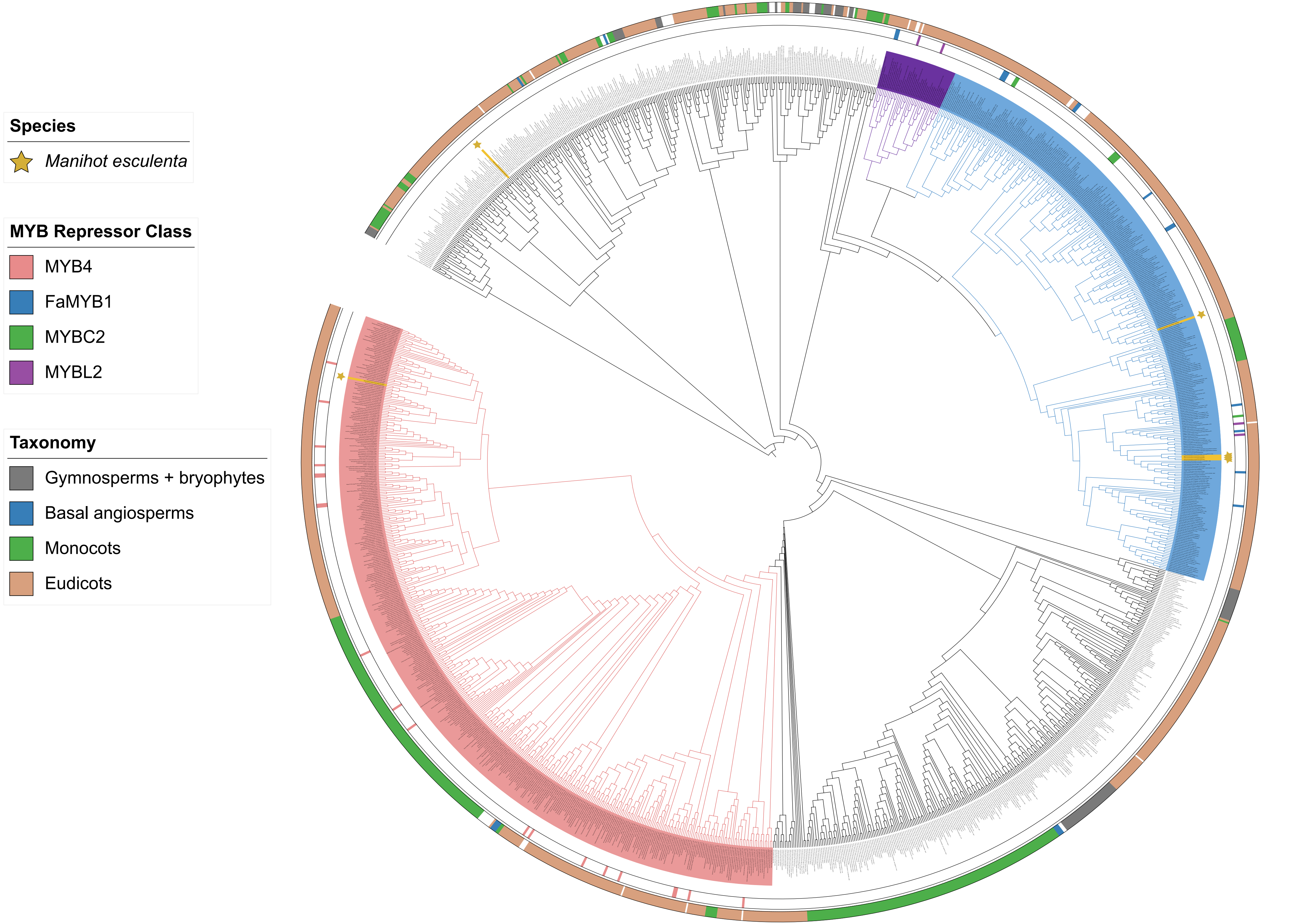

### Additional File I

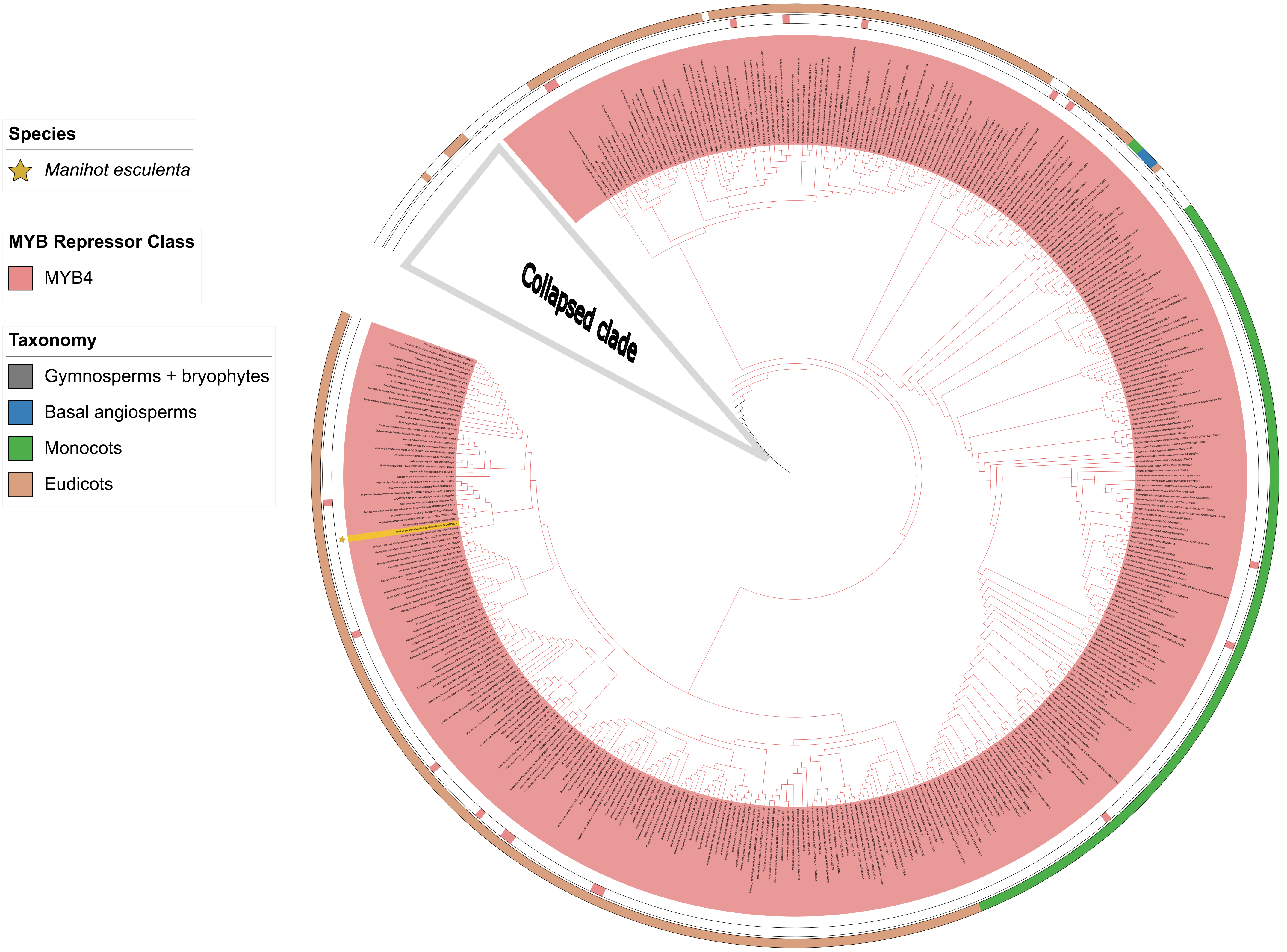

### Additional File J

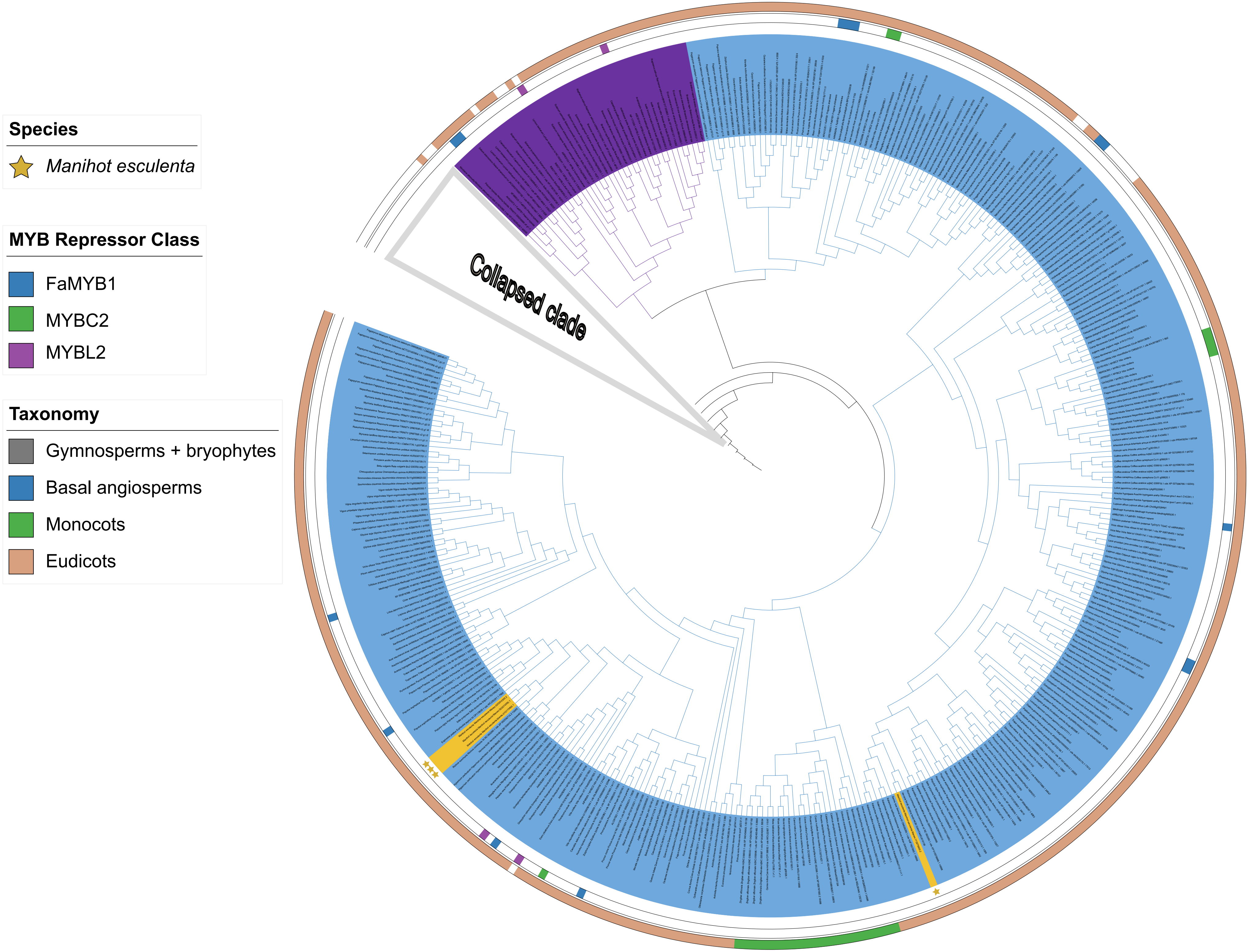
